# Natural versus Random Proteins: Nouvel Neural Network Approach Based on Time Series Analysis

**DOI:** 10.1101/687558

**Authors:** Alexei Tsygvintsev

## Abstract

We study the set of about 35000 primary structures of natural proteins of length more than 360 residues and the same size set generated via partial or total randomization. Associated to every sequence composed of 20 amino acids, a time series is formed from hydropathy values of the first 360 residues. To measure the absolute deviations of hydropathy index on different time scales, the 24-dimensional vector of total log-amplitudes is introduced. We describe then a configuration of the 1-hidden layer neural network which is trained to solve the binary classification problem of natural and random sequences. A satisfactory distinguishing accuracy random/natural of 88% is obtained.

## 1. Introduction

The term Never Born Proteins was originally introduced in [1] to describe a collection of randomly generated sequences composed of 20 amino acids which could posses some folding stability properties observed in the laboratory experiments. It seems nevertheless, that from the purely combinatoric point of view, there is no clear difference between primary structures of natural proteins observed in Nature and synthetic amino acids sequences randomly uniformly generated [9]. Based on the Shannon entropy [7], it was reported in [6] that natural protein sequences are random like Never Born Protein sequences i.e their corresponding entropies are asymptotically same. It should be noted that in the above and similar studies only primary structures, viewed as words composed of 20 amino acids, are considered. By contrast, many authors have stressed the importance of the quantitative features emerging from particular secondary or tertiary structures of proteins (experimentally known for natural and predicted for random ones) with the use of which more sharp distinction between random and natural can be achieved (see for example [8] where the evolutional network was designed for this purpose). At the same time, a considerable number of controversial viewpoints on the question of random/natural proteins can be found in literature and apparently many conclusions are methodologically driven. A number of aspects of this problem require further investigation and clear precise definitions.

In this communication we address the problem of sorting out random/natural applying the artificial neural network approach based solely on primary structures and hydropathy values of individual amino acids. We study an initial data base of 35022 natural proteins of length 361 ≤ *N* ≤ 400 residues taken from UniProtKB (http://www.uniprot.org) and the same size data set formed by uniformly randomised (at different degrees) sequences of the same length.

The particular size of *n* = 360 residues is chosen by considering a particularly high number of divisors of *n* which is 24.

We define in Section 2 the 24-dimensional vector, associated to every amino acid sequence *S* (natural or randomised) of the length *n*, whose entries are logarithms of the sum of local amplitudes computed for every partition of *S*. The corresponding time series is constructed in a natural way by adding the *z*-score values of hydropathy parameters [5] of amino acids along the sequence *S* starting from the first left residue. See [4] for some alternative applications of discrete time series representations of protein sequences.

In Section 3, a conventional in machine learning partition 80 % - 10 % - 10 % of the initial data set to training-validation-test sets is applied. We use the 24 - 24 - 1 neural network trained to solve the binary classification problem random/natural for different length of randomized tails of natural primary structures. The trained neural network is capable to classify correctly 85 % of the testing set containing 50 % of random and 50 % of natural amino acids chains.

## 2. Total amplitudes of amino acid sequences

To analyse the primary structure of proteins, in order to use the time series tools, we need to transform a given sequence of amino acids, composed of 20 residues {*A, C, D, …*}. into the numerical form. We begin by considering the hydropathy values of amino acids, defined in [5] (Table 1) which are then normalised using *z*-score (Table 2).

**Table 1.** Hydropathy values of 20 amino acids

| A | C | D | E | F | G | H | I | K | L | M | N | P | Q | R | S | T | V | W | Y |
| --- | --- | --- | --- | --- | --- | --- | --- | --- | --- | --- | --- | --- | --- | --- | --- | --- | --- | --- | --- |
| 1.8 | 2.5 | -3.5 | -3.5 | 2.8 | -0.4 | -3.2 | 4.5 | -3.9 | 3.8 | 1.9 | -3.5 | -1.6 | -3.5 | -4.5 | -0.8 | -0.7 | 4.2 | -0.9 | -1.3 |

**Table 2.** *Z*-score Hydropathy values

| A | C | D | E | F | G | H | I | K | L | M | N | P | Q | R | S | T | V | W | Y |
| --- | --- | --- | --- | --- | --- | --- | --- | --- | --- | --- | --- | --- | --- | --- | --- | --- | --- | --- | --- |
| $H_1$ | $H_2$ | $H_3$ | $H_4$ | $H_5$ | $H_6$ | $H_7$ | $H_8$ | $H_9$ | $H_{10}$ | $H_{11}$ | $H_{12}$ | $H_{13}$ | $H_{14}$ | $H_{15}$ | $H_{16}$ | $H_{17}$ | $H_{18}$ | $H_{19}$ | $H_{20}$ |
| 0.7 | 1.0 | -1.0 | -1.0 | 1.1 | 0.0 | -0.9 | 1.6 | -1.1 | 1.4 | 0.8 | -1.0 | -0.3 | -1.0 | -1.3 | -0.1 | -0.0 | 1.5 | -0.1 | -0.2 |

For the sake of simplicity, we provide the truncated values only while 16-digits precision is used in all numerical computations.

Let *S* = (*S*_1_, *S*_2_, *…, S*_360_), *S*_*i*_ ∈ Σ be any sequence of amino acids where

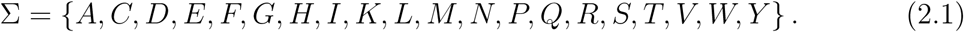

The time series *X* associated to *S* is defined as follows

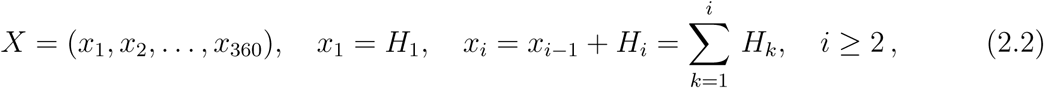

by adding recursively the *z*-score hydropathy values along the amino acid chain starting from its first residue on the left. Figures 1–4 contain graphs of these series for 4 particular natural proteins. We observe that their structures and regularity can be quite different : from the sideways behaviour (Fig.1–2) to the trend-like one (Fig. 3–4). Fig. 5 illustrates the time series of a typical synthetic sequence whose all residues (excepting the very first one which is fixed to be “M”) are chosen randomly.

**Figure 1.**
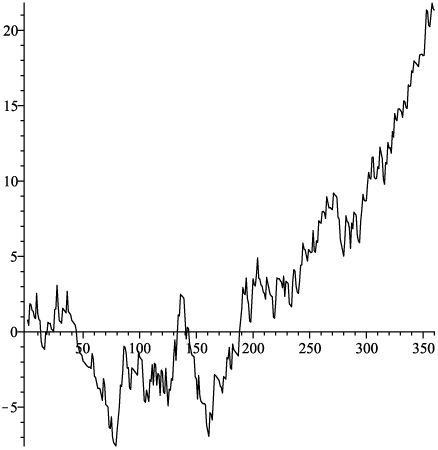
Time series formed by first 360 residues of of the Putative movement protein, Q91TW8.

**Figure 2.**
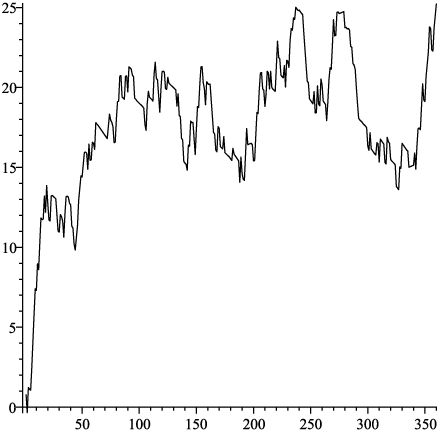
Probable GPI-anchored adhesin-like protein PGA32, Q5ADQ7.

**Figure 3.**
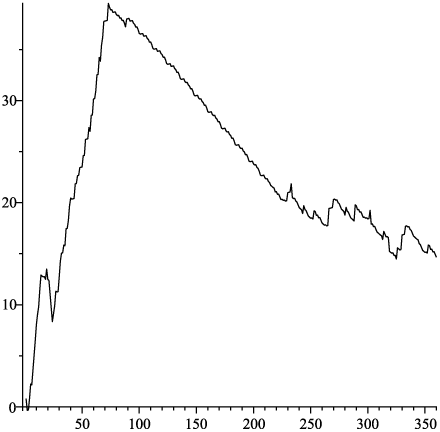
Prisilkin-39, C0J7L8.

**Figure 4.**
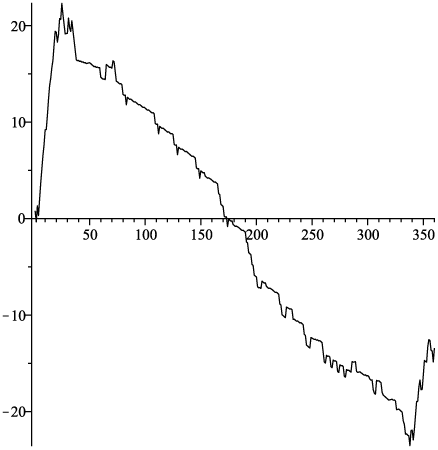
29C-likeproteinDDB G0287399, Q54KD5.

**Figure 5.**
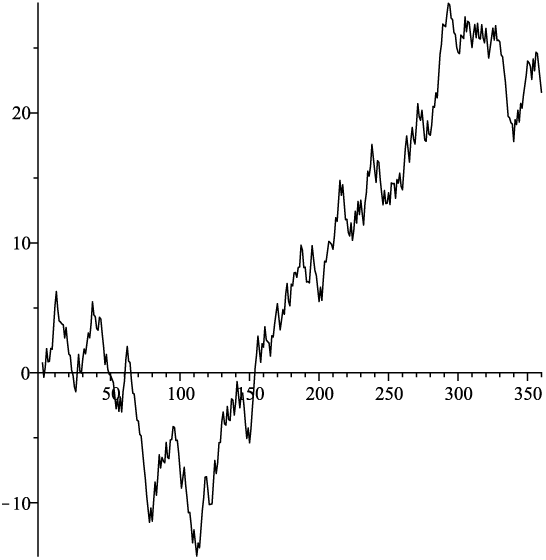
Uniformly random sequence of 360 residues.

We fix now *n* = 360 and define

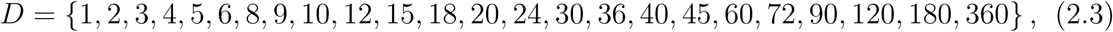

the set of all 24 divisors of *n*.

For a particular divisor *d* ∈ *D* one gets a partition of *X*

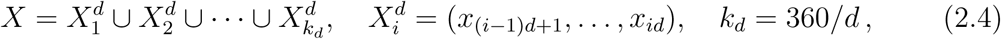

in *k*_*d*_ fragments of equal size *d*.

For *i* ∈ {1, *…, k*_*d*_}, the corresponding local *i*^th^ amplitude is defined according to

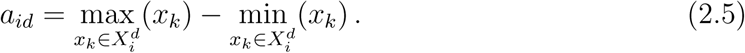

The sequence of sums of local amplitudes

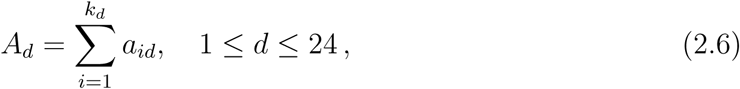

is mapped then to the 24-dimensional vector

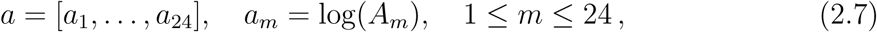

called the *total log-amplitude* of the time series *X*. Figures 6–7 contain graphical representations of *a* for two particular natural and random truncated primary sequences. The linear and correlated patterns can be clearly spotted. This is not really surprising, since, as was reported in [2], for many stochastic time series, some entries of total log-amplitude vectors (2.7) obey linear law with the slope given by the fractal dimension of the series in question.

**Figure 6.**
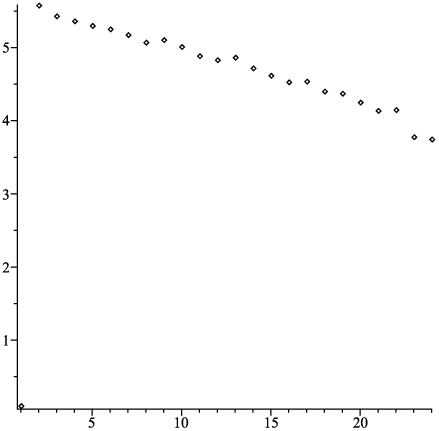
The total log-amplitude vector for random sequence from Fig.5.

**Figure 7.**
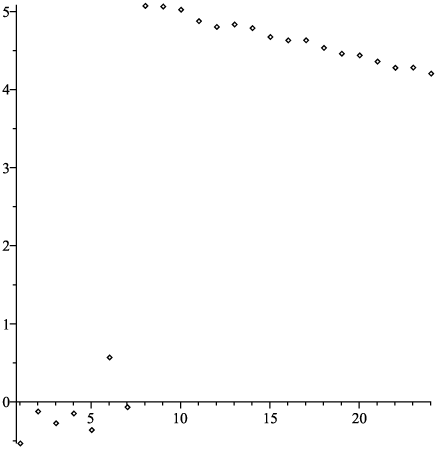
The total amplitude vector of 360-residues left tail of GDSLesterase, Q9SVU5.

## 3. Neural network description and solving the classification problem

For numerical simulations, the Java neural network framework Neuroph 2.96 was used.

First, we selected the data set DATA from UniProtKB (http://www.uniprot.org) of primary structures of natural proteins of the length 361 ≤ *N* ≤ 400 which contains 35022 sequences (rows) in total. This set was systematically row shuffled before each new network training. Then, DATA is partitioned into 3 subsets according to

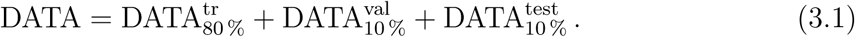

For every of the 3 above subsets, its *α* %-randomised “clone” is created to form a parallel to (3.1) randomised partitioning

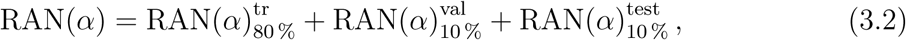

where every row of the table 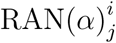 is obtained by uniform randomization 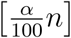, *n* = 360 amino acids of every row of the table 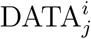 starting from the last residue on the right, while leaving others to be intact (with the very first residue on the left “M” always kept unchanged).

The sets of both types are then joined together to build a new data set

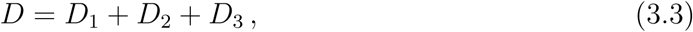

with

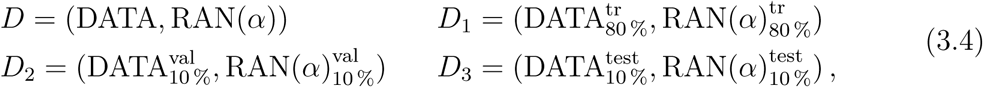

which is used to train the 24 - 24 - 1 neural network. The Sigmoid activation function was chosen and the *z*-score normalisation was applied to input vectors whith parameters (empirical average and variance) computed from the 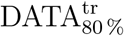 set only. The desired values of the network’s data set were fixed according to: “0” ~ *α* %-random input row and “1” ~ natural primary structure row. Training was done using the back-propagation in a batch mode using the *D*_1_ set. To define the stoping rule, one waits until the LMS error computed on the validation *D*_2_ set starts to increase while the LMS error evaluated on the training *D*_1_ set is still decreasing.

## 4. Results and conclusion

The capacity of trained network was evaluated to classify sequences from the test set *D*_3_, not used during the training, and which composition was 50*/*50 natural or random. The results are quite satisfactory and listed in Table 3.

**Table 3.**
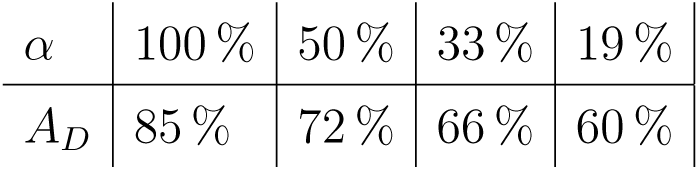
Distinguishing accuracy *A*_*D*_ of the neural network for different *α*’s

Decreasing of *A*_*D*_ with lower values of *α* is expectable since *A*_*D*_ → 50% as *α* → 0 with the difference random/natural disappearing. The maximum 88% of the accuracy can be explained by the fact that only limited part of protein’s structure is coded by hydropathy values of amino acids. Our preliminary findings indicate that neural networks are able to uncover hidden distinguishing patterns in total log-amplitude vectors of natural and random amino acid sequences. Since these numerical features are intimately correlated to fractal and self-similarity characteristics of time series [2], it would be of interest to establish links between our results and other studies of the protein fractal structures [3].

## Acknowledgments

The author is grateful to I. Gannaz end C. Combet for fruitful discussions and provided valuable assistance in accessing the protein data base.

